# Behavioral response of insecticide-resistant mosquitoes against spatial repellent: a modified self-propelled particle model simulation

**DOI:** 10.1101/2020.03.20.000281

**Authors:** Guofa Zhou, Leonard Yu, Xiaoming Wang, Daibin Zhong, Ming-chieh Lee, Solomon Kibret, Guiyun Yan

## Abstract

Rapidly increasing pyrethroid insecticide resistance and changes in vector biting and resting behavior pose serious challenges in malaria control. Mosquito repellents, especially spatial repellents, have received much attention from industry. Many of these repellents contain the same or similar chemicals as those used in insecticides. Does resistance to insecticides affect the efficacy of spatial repellents? We attempted to simulate interactions between mosquitoes and repellents using a machine learning method, the self-propelled particle (SPP) model, which we modified to include attractiveness/repellency effects. We simulated a random walk scenario and scenarios with insecticide susceptible/resistant mosquitoes against repellent alone and against repellent plus attractant (to mimic a human host). We assumed attractant odors and repellent chemicals diffused randomly and omnidirectionally, and that mosquitoes were confined in a cubic cage. We modified the velocity and direction components of SPP using attraction/repulsion rates and concentrations. Simulation results indicated that without attractant/repellent, mosquitoes would fly anywhere in the cage at random. With attractant, mosquitoes might or might not fly toward the source (i.e., the human), depending on the simulation time (and thus the attractant concentration distribution). Eventually, however, all mosquitoes were attracted to the source of the odor. With repellent, results depended on the amount of chemical used and the level of mosquito insecticide resistance. All insecticide-susceptible mosquitoes eventually moved to the corner of the cage farthest from the repellent release point. Surprisingly, a high proportion of highly resistant mosquitoes might reach the attractant release point (the human) earlier in the simulation when repellent was present compare to no repellent was present. At fixed concentration, a high proportion of mosquitoes could be able to reach the host when the relative repellency efficacy (compare to attractant efficacy) was <1, whereas, no mosquitoes reached the host when the relative repellency efficacy was > 1. This result implies that repellent may not be sufficient against highly physiologically insecticide resistant mosquitoes, since very high concentrations of repellent are neither practically feasible nor cost-effective.

## Background

Malaria remains to be the world’s most common mosquito-borne disease, with an estimated 228 million cases worldwide in 2018 [1]. Control efforts mainly rely on vector control using a single class of insecticides, the pyrethroids, which is the only class approved for use on long-lasting insecticidal nets (LLINs) [2]. Pyrethroids, along with other pesticides, are also widely used to control agricultural pests on livestock and field crops worldwide [3]. The past decade has seen a dramatic increase in reports of physiological pyrethroid resistance in malaria vectors [4–15]. The rapid increase in pyrethroid resistance necessitates an immediate proactive resistance management response to avoid compromising existing effective interventions. In addition to the increase and spread of insecticide resistance, the biting and resting behaviors of malaria vectors have evolved. A number of recent studies have documented changes in the biting behavior of the major African malaria vectors, *Anopheles gambiae* and *Anopheles funestus*, from midnight biting to biting in the early evening and morning hours, and to biting outdoors, where people are not protected by indoor residual spraying (IRS) or LLINs [16–18]. This early outdoor biting behavior is likely due to insecticide-induced behavioral changes in malaria vectors, i.e., avoiding contact with insecticide-treated bed nets and walls [19, 20]. Residual malaria transmission has become a very important challenge in malaria control [21].

Various alternative vector control tools against outdoor transmission have been the subject of research studies in recent years [22–25]. Among these tools, repellents have received the most attention from industry. The global mosquito repellent market was valued at approximately $3.2 billion in 2016 and is expected to reach ∼$5 billion in 2022, growth driven chiefly by recent outbreaks of mosquito-borne diseases [26]. Repellents come in two types: topical and spatial. Topical repellents such as DEET require constant application and reapplication on skin, clothing, or other surfaces to discourage mosquitoes (and arthropods in general) from landing or climbing on the surface [27–34]. Spatial repellents release into the air volatile active ingredients that interfere with mosquitoes’ ability to find a host, thus preventing mosquitoes from contacting the host and taking a blood meal, thus preventing disease transmission [35–40]. Spatial repellents confer protection against mosquito bites through the action of emanated vapor or airborne chemical particles in a large space, such as a room or a yard surrounding a household [35–42]. Due to their relatively low cost and ease of deployment, spatial repellents are popular in developing regions [43]. Spatial repellents may significantly aid in preventing mosquito-borne diseases if properly incorporated into integrated vector management approaches [40]. The best known spatial repellent is the burning coil, which releases the chemical into a space such as a room, preventing mosquitoes from entering the entire space [43].

However, spatial repellents, such as some coil products, use pyrethroids and other synthetic chemicals and plant products as major volatile ingredients to repel mosquitoes [23, 25, 44–46]. These volatile chemicals prevent mosquitoes from feeding on humans through several mechanisms, including 1) knockdown and mortality; 2) deterrence (mosquitoes are prevented from entering human dwellings); 3) irritancy (mosquitoes enter houses but leave early); 4) excito-repellency (mosquitoes exit the house to avoid contacting airborne insecticides); and 5) feeding inhibition (mosquitoes are prevented from biting and getting blood meals) [23, 25, 35, 37, 40, 44–50]. Since malaria vectors have developed widespread resistance to insecticides, the efficacy of spatial repellents against these resistant mosquitoes is unknown. Some recent studies have found that insecticide-resistant mosquitoes behave differently than susceptible mosquitoes when they encounter repellents. For example, Deletre et al. found that DEET had a reduced repellency effect against resistant strains of *An. gambiae* but maintained irritancy for the susceptible strain [51]. Similarly, Agossa et al. found that as a result of resistance, pyrethroid-based malaria control tools have decreased toxicity and repellent effects [52]. Kawada et al. found that the frequency of takeoffs from a pyrethroid-treated surface and flying times without contacting the surface increased significantly in pyrethroid-susceptible *An. gambiae* s.s., while the *An. gambiae* s.s. wild colony (i.e., the resistant strain) exhibited no such behavior [53]. Studies of *Anopheles* and *Aedes* mosquitoes show similar results, i.e., physiological insecticide resistance modifies vector contact avoidance behavior, and that, in general, a higher concentration of pyrethroids is needed to deter blood-feeding by resistant vectors [54–58]. While the evidence from these studies points to that repellents have a reduced repellency effect against physiologically insecticide resistant mosquitoes, it is not conclusive. In addition, data are limited on the response to spatial repellents by mosquito populations that have evolved outdoor biting behavior.

Machine learning methods such as self-propelled particle (SPP) models have offered innovative ways to study the collective behaviors of living organisms and their interactions with their environments, including colonies of wasps, schools of fish, flocks of birds, and others [59–63]. Many animals can be treated as SPPs; they find energy in their food and exhibit various locomotion strategies, from flying to crawling. These biological systems can propel themselves based on the presence of chemoattractants [63]. A number of SPP models have been proposed, ranging from the simplest Active Brownian Particle model to highly elaborate and specialized models aimed at describing specific systems and situations [64, 65]. For example, an SPP model simulation found that as locust population density increased, locusts changed their behavior from relatively disordered, independent movement of individuals within the group to the group moving as a highly aligned whole [65]. This modeling result was supported in the field: when locust density reached a critical value of 74 locusts/m^2^, the locusts ceased making rapid and spontaneous changes of direction and instead marched steadily in the same direction for the full 8 hours of the experiment [66]. *Anopheles* mosquitoes, which find their hosts via human odors, can be treated as SPPs because they propel themselves based on chemoattractants [67, 68]. Mosquito host-seeking involves the movements of individual females as well as the collective behaviors of groups that result from individuals’ local interactions with each other and with their environment [67, 68]. To model mosquito reactions to spatial repellents, however, other parameters must be considered in the presence of repellents, because it is not clear how insecticide-resistant mosquitoes respond to these repellents. These additional parameters may be incorporated by modifying existing models, since SPP models allow for particle assemblies to be temporally and spatially reversible in complex media, e.g., media that includes barriers resembling a bed net or repellent situation faced by mosquitoes [69–76].

The aim of this study is to demonstrate mosquitoes’ host-seeking behavior in the presence of spatial repellent with/without resistance using machine learning simulations, and to examine how physiological insecticide resistance impacts this behavior at the individual and population level. The specific question asked is: Can physiologically insecticide-resistant mosquitoes reach the host before being repelled by the spatial repellent? If the answer is yes, what proportion can reach the host before they are repelled? Such knowledge is crucial in developing more efficient methods which improve the field effectiveness of spatial repellents and other means of mosquito control.

## Methods

### Model development

In this study, we considered the effects of two elements: i) attractive odors to mimic human odor and attract mosquitoes, and ii) spatial repellents to repel mosquitoes and protect humans from mosquito biting. Mosquito flight behavior was simulated using the constrained SPP model [70, 73, 74]. We chose this approach based on the prior success of SPP in modeling insect social behaviors [69–75]. We modified the SPP model by adding attractant and repellent to influence the mosquito flight path. We used both insecticide-susceptible and -resistant mosquitoes, since insecticide-resistant mosquitoes may behave differently than susceptible mosquitoes when they encounter spatial repellent. Mosquitoes were thus able to choose their respective paths and move non-uniformly into the spatial repellent–affected space. This constrained, non-uniform movement helped insecticide-resistant mosquitoes to avoid the repellent.

Our modified SPP model, based on the Vicsek model [77], is shown in equation (Eq. 1). In this model, mosquito flight velocity, and location, is simulated as a function of Brownian motion and modulated by attractants and repellents [78]. With neither attractant nor repellent, we assume mosquitoes move randomly. With attractant, mosquitoes are drawn toward the attractant release point; conversely, mosquitoes are driven away from the repellent release point. The flying velocity of a mosquito at a given time is modeled as:

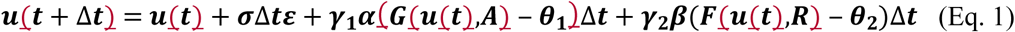

where *u*(*t*) is a 3-dimensional vector and parameters are as shown in Table 1. To simplify the simulation process, we assume mosquitoes are confined within insect-proof cages. Mosquitoes cannot escape, they have no food source other than the attractant odor, and the repellent is pure, with no bed net or IRS use. These conditions rule out any potential outdoor resting or other avoidance behavior.

**Table 1.** Descriptions of parameters used in the models

| Parameter | Description | Initial value |
| --- | --- | --- |
| $\Delta t$ | Time and time interval for each step of simulation | 0.03 s |
| $\sigma$ | Random walk rate in the form of Brownian motion | 1.0 |
| $\varepsilon$ | Gaussian random variable with mean of 0 and variance of 1 | |
| $\gamma_1, \gamma_2$ | Mosquito response rates | 1.0, 1.0 |
| $\alpha, \beta$ | Attractance/repellency efficiency | 0.5, 0.5 |
| $\theta_1, \theta_2$ | Attractant/repellent release locations | |
| $\rho$ | Response scale parameter | |
| $a$ | Response decay/increase rate parameter | 1.0 |
| F, G | Attractant/repellent response functions in the form of K with different $a$ and $\rho$ | |
| A, R | Attractant/repellent concentrations in the form of C with different D, $\mu$ and $C_0$ | 5.0, 5.0 |
| $C_0$ | Attractant/repellent release rate | |
| D | Diffusion coefficient | 1.0 |
| $\mu$ | Scale parameter | 1.0 |

The mosquitoes’ response to attractant/repellent depends on the concentration of attractant/repellent and the mosquitoes’ response strength; i.e., insecticide-resistant and -sensitive mosquitoes respond differently to repellent. The mosquito response to attractant/repellent is modeled as an inverse-exponential-decay function:

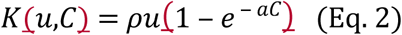

where *C* is the attractant/repellent concentration and *a* is the response parameter. *C* is a function of space and time and is modeled as a convection-diffusion model with a point source [79]:

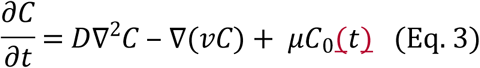

where *C*_0_ is the attractant/repellent release rate, *v* is the velocity field that the quantity is moving with (e.g., wind or temperature), D is the diffusion coefficient, μ is the scale parameter which measures the attractant/repellent releasing strength, and ∇ represents the concentration gradient.

An attractant/repellent’s concentration is affected by its release rate. For simplicity, we assume it is not contingent on temperature, wind speed (i.e., *v* = 0), or delivery mechanism. In an individual household setting in rural Africa, a repellent such as burning coils usually can be seen as a point source with a constant rate of release, whereas a real person releasing human odor is not really a point source. However, for simplicity of simulation, we assume both attractant and repellent are released from a point source with constant speed of release.

### Mosquito movement rules

In our simulations, an individual mosquito was treated as point particle; body size was omitted. Mosquitoes had no effect on each other during movement, either attractive or repulsive. Mosquito motion was restricted to the ‘sealed’ house. Both the repellent volatile chemicals and the attractant odors had threshold concentrations, below which the mosquitoes navigated randomly; i.e., attractant/repellent did not affect mosquito movement when odor/volatile chemical concentration was low. For concentrations above the threshold, mosquitoes responded by changing their moving behavior, moving away from higher levels of repellent or toward the odor release point. High concentration levels biased mosquitoes’ flight direction, resulting in flights that were on average moving away from the repellent or toward the attractant. However, we assumed that repellent/attractant concentration did not affect mosquito flight speed; i.e., average speed and deviation did not change.

Step length for mosquito movement was drawn separately for X/Y/Z directions from Brownian motion, with a mean of 1 cm and variance of 1 cm^2^. Based on video tracking of flying speed of *Anopheles arabiensis* and *Anopheles gambiae* sensu stricto against untreated nets and LLINs [68, 80], we assumed mosquitoes move 1000 steps per 30 s, which was equivalent to average of 10 m per 30 s, same as observed in the video tracking experiments. Direction of movement was determined by (X-1.0, Y-1.0, Z-1.0). However, directions were adjusted either toward the attractant release point or against the repellent release point, depending on the resistance level and media concentration.

### Host odor and repellent

Host odor and repellent were released independently at different emission and diffusion rates, both following the random diffusion principle as described in (Eq. 3). For simplicity we assumed no other media affected the diffusion of attractant and repellent. However, we did vary the attractiveness and repellency rates. There were no other constraints, so both attractant and repellent could potentially fill the entire experimental space, but we did set a maximum concentration level within the space. (In the real world, houses are not sealed and odor and repellent chemicals can diffuse out of the house, which means indoor concentration remains relatively stable rather than increasing indefinitely.) As stated above, we assigned a threshold concentration for both repellent and attractant. Below the threshold, mosquitoes navigated randomly. Above the threshold, mosquitoes moved away from the repellent move toward the host odor.

### Mosquito resistance

Because resistant mosquitoes tolerate higher insecticide concentrations, we assumed they also tolerate a higher concentration of the volatile chemicals released into the air by the repellent. In other words, host-searching movements of insecticide-resistant mosquitoes may not be inhibited by repellent if the concentration of volatile chemicals is low. We made this assumption based on observations that spatial repellent had a delayed impact or decreased toxicity and repellent effects against field-caught insecticide-resistant mosquitoes [52, 55–58, 81]. For this study, we assumed resistant mosquitoes tolerated a 2-fold higher concentration of repellent compared to non-resistant/susceptible mosquitoes.

### Additional rules

We assumed mosquitoes did not rest on the cage walls, as if the mosquitoes were starving when released into a room with a sleeping host and would not stop flying until they reached the host. Mosquitoes could not escape from the cages. For simplicity of simulation we also assumed no knockdown or mortality effects; this is probably not accurate, since many spatial repellents consist of pyrethroid insecticides (although it is accurate if the concentration of volatile chemicals is low).

### Outcome measurements

We measured the following parameters: average landing time, landing rate, average repelling time, repelling rate, distribution of landing rate, and distribution of repelling rate. Landing on a human or host was defined as a mosquito moving to <1 cm (or 1 step) from the attractant odor release point. A mosquito was repelled if it was pushed out of the enclosed space, i.e., moved to <1 cm from the corner of the cage farthest from the repellent release point.

In real-world hut experiments, three indicators are used to measure repellent efficacy: deterrence, excito-repellency, and toxicity [82, 83]. Deterrence is determined by comparing the total number of mosquitoes in cages with spatial repellent to the number in control cages without spatial repellent. For the simulation study, we removed a mosquito once it reached the attractant (found the blood source) or was expelled to the far end of the cage opposite the repellent release location (exited the house). Excito-repellence is measured as the proportion of mosquitoes found at the far end of the cage opposite the repellent release location, in the spatial repellent treatment relative to the control. Toxicity is determined by comparing the mortality rate in spatial repellent treatment houses to that of the control houses. In this study, we assumed repellent only repels but does not kill mosquitoes.

### Simulation of mosquito response to repellent with/without attractant

Simulations were conducted under four scenarios: a) random walk, i.e., no attractant and no repellent; b) random walk modified by adding attractant to mimic human host; c) random walk modified by adding repellent to simulate prevention efficacy; and d) random walk modified by adding both attractant and repellent. We also simulated different mosquito resistance levels: sensitive (no resistance at all) and resistant (insensitive to a 2-fold higher level of repellent) [51, 52]. Resistance levels were simulated by reducing the repellency rate [51–53].

Our simulations assumed mosquitoes were released in the middle of an enclosed space with a size of 5m × 5m × 3m, which similar to the size of typical African house [68]. Both attractant and repellent were released on the ground in one corner of the space, and both were treated as point objects. In each simulation, 10 mosquitoes were released for up to a maximum of 30 minutes. The simulation was repeated 100 times for each scenario with both susceptible and resistant mosquitoes.

Simulations were realized using R 3.5.2.

## Results

### Flight path tracking

Figure 1 illustrates the potential mosquito flight paths (n = 10 mosquitoes) simulated by the model under different settings: random walk without attractant or repellent, with attractant, and with repellent (see Supplement Figure S1 for single mosquito flying path). By 30 seconds, 3 of the 10 mosquitoes have been attracted to the “human,” i.e., the attractant release point (Figure 1B, middle). Figure 2 shows the 2D projected attractant/repellent concentration and the locations of all mosquitoes at different points in time. Here, simulations indicated that all non-resistant mosquitoes were repelled to the corner farthest from the repellent release point within 1 minute (Figure 2C), whereas most of the resistant mosquitoes were able to stay in areas with low repellent concentration (Figure 2D). The presence of repellent did make mosquitoes more difficult to find the host even if when repellent concentration was low (Supplement Figures S1.B & S1.D).

**Figure 1.**
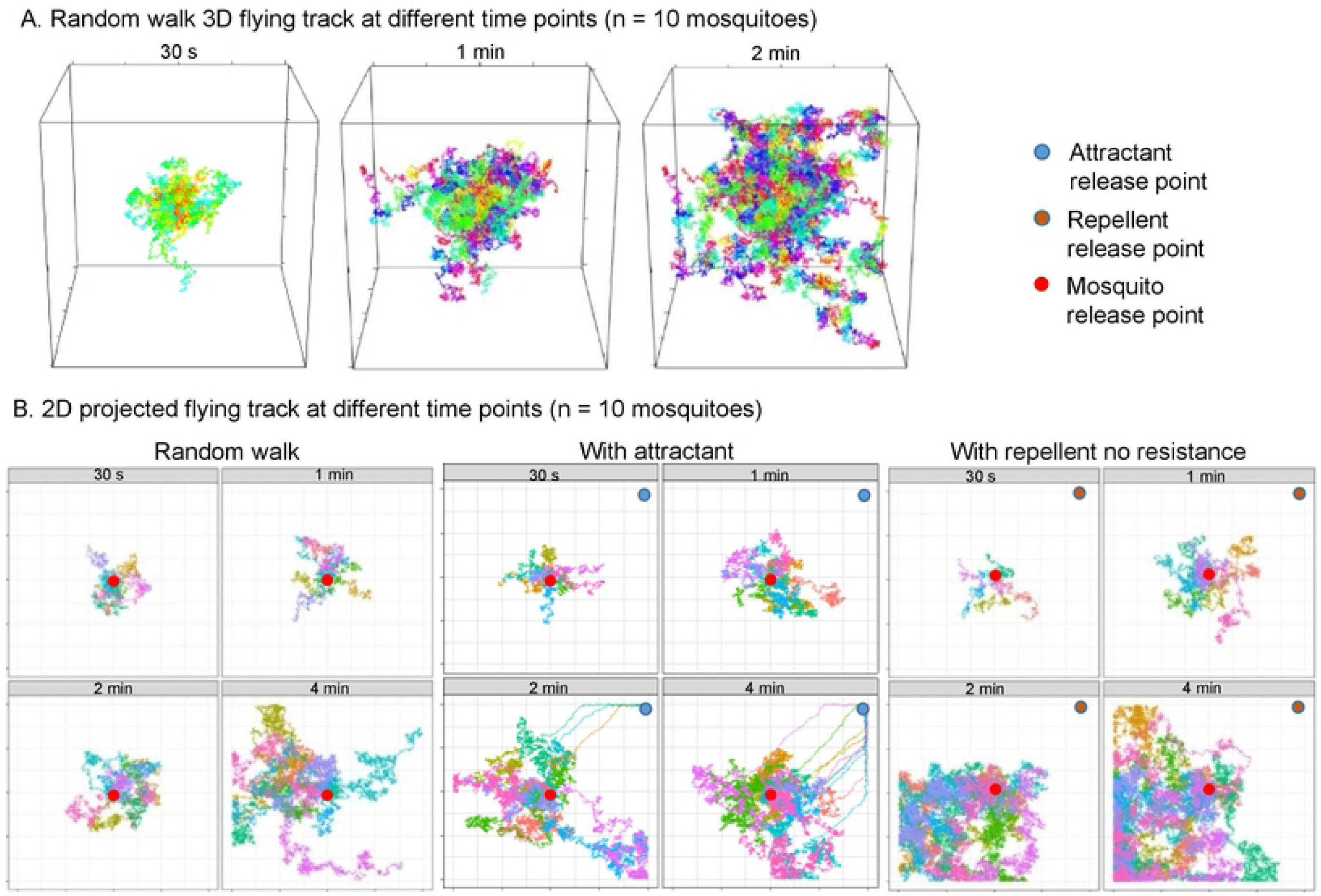
Illustrations of simulated mosquito flying paths. A) 3D display of random walk at different time points; B) 2D projected flying paths at different time points for random walk (left panel), with attractant (middle panel), and with repellent and no resistance (right panel).

**Figure 2.**
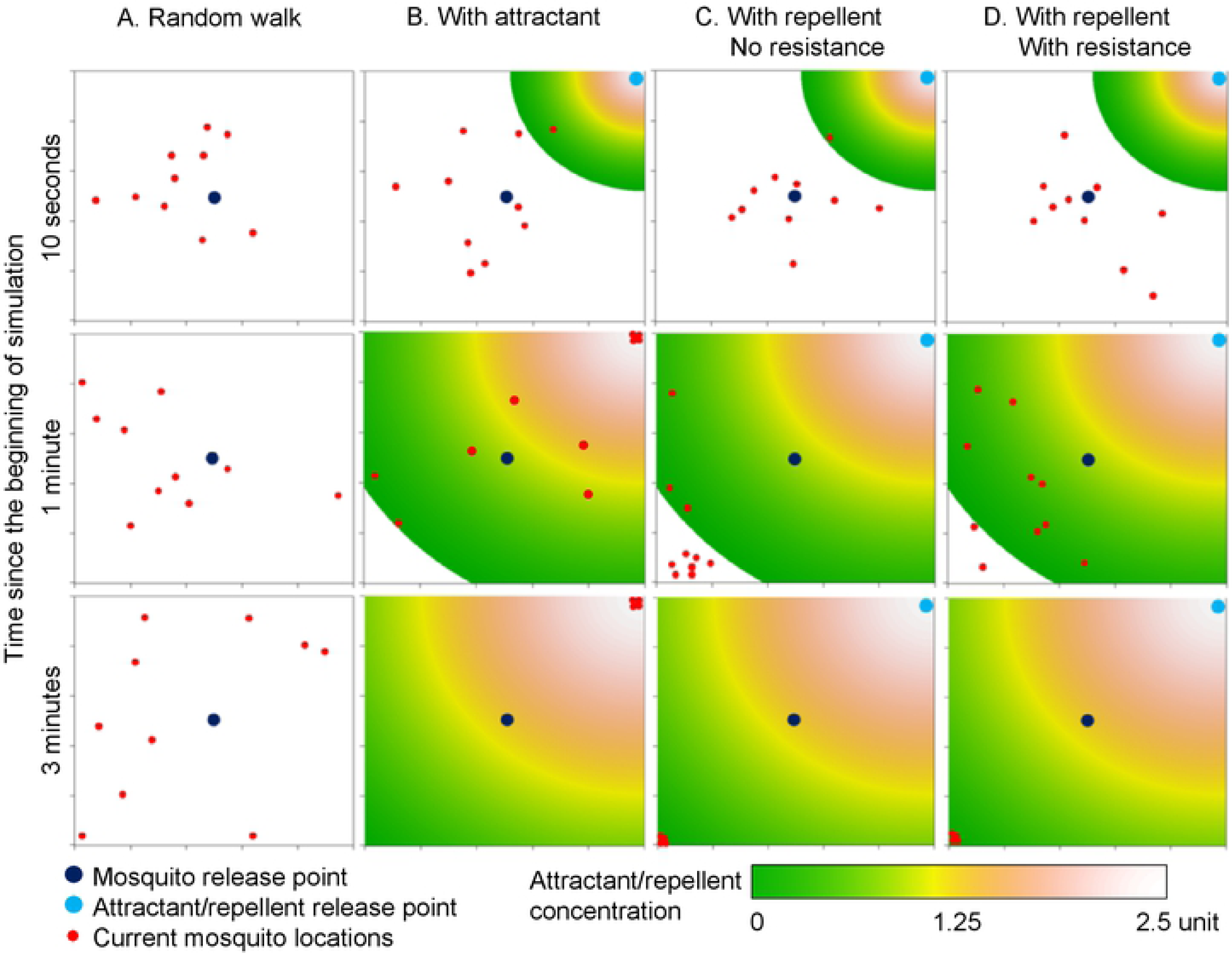
Current 2D projected locations of mosquitoes at different time points in different settings. A) random walk; B) with attractant; C) with repellent and no resistance; and D) with repellent and resistant mosquitoes. Colors in the heat maps show different concentrations of attractant odors or repellent volatile particles.

### Attractant effect

Figure 3 shows how attractant concentration affected mosquito host-searching behavior. In low-concentration settings (C_0_ = 5 unit), the first mosquito had reached the host by 38 s; about 50% of the mosquitoes had reached the host by about 45 s; and all mosquitoes had reached the host by 117 s (Figure 3). The average time to reach the host was 83.1 (SD 1.5) s.

**Figure 3.**
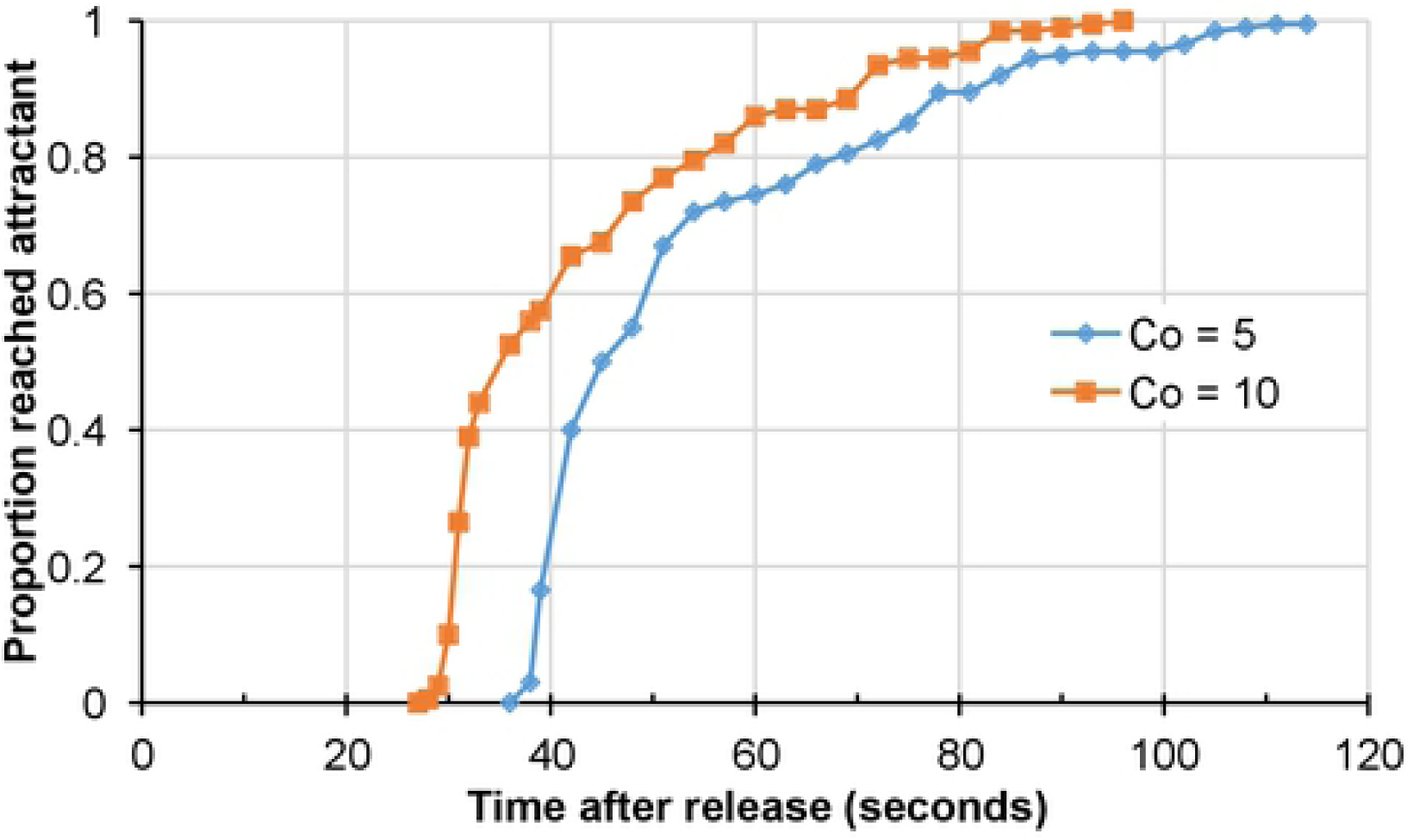
Frequency distribution of time when mosquitoes reached the host at different attractant release rates. C_0_ is the attractant concentration.

If the attractant release rate was doubled (C_0_ = 10 unit), the first mosquito reached the host by 28 s and all mosquitoes reached the host by 96 s (Figure 3). The average time to reach the host was 65.7 ± 1.5 s.

### Repellent effect

Figure 4 shows the distribution of times at which mosquitoes were repelled; no attractant was released in these simulations. For the non-resistant mosquitoes, some mosquitoes were repelled as early as 170 s after the repellent was first released, and all were repelled by 198 s (Figure 4). The average repelling time was 194.5 ± 27.3 s. The resistant mosquitoes started being repelled at 258 s, and all were repelled by 279 s (Figure 4). The average repelling time was 275.5 ± 42.6 s.

**Figure 4.**
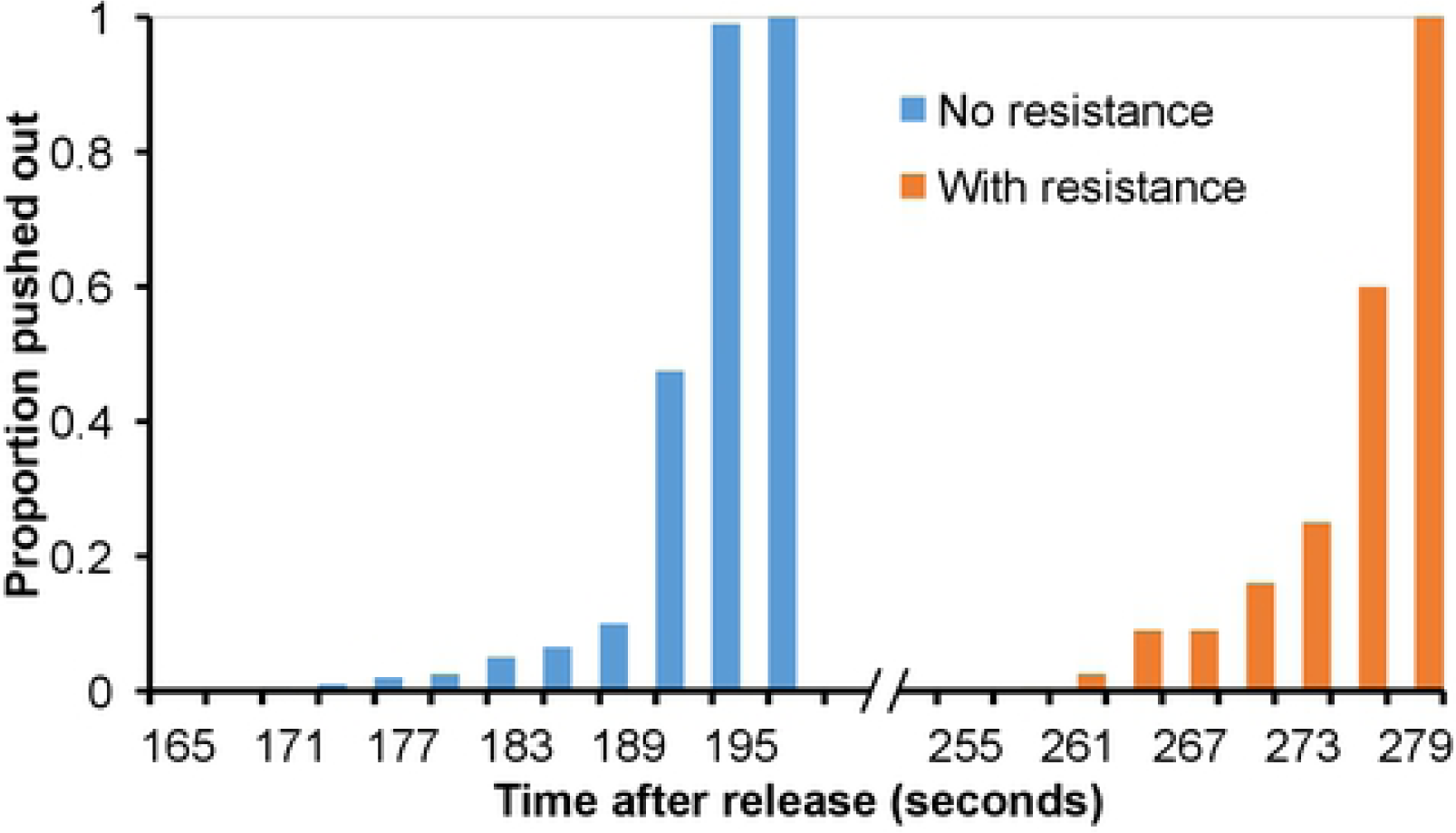
Frequency distribution of time when mosquitoes were expelled, for non-resistant (blue bar) and resistant (red bar) mosquitoes.

### Attractant plus repellent

Here we assumed that the attractant odor and the repellent were released starting at the same time. For the resistant mosquitoes, relative repellency efficiency (β/α) played an important role when maximum repellent/attractant concentrations were fixed (Figure 5). If repellency efficiency was low (β/α = 0.8), the simulation results indicated that a surprisingly very high proportion, up to 97%, were able to reach the host before they were repelled (Figure 5A). On average these mosquitoes reached the host within 40.7 ± 4.2 s (range 32–48 s), after which they were pushed away from the host by the increased repellency. However, they were not pushed out of the simulation space until 255 s (Figure 5A). Eventually, all mosquitoes were repelled, with an average repel time of 278.1 ± 37.7 s, and all mosquitoes were repelled within about 30 s (Figure 5A).

**Figure 5.**
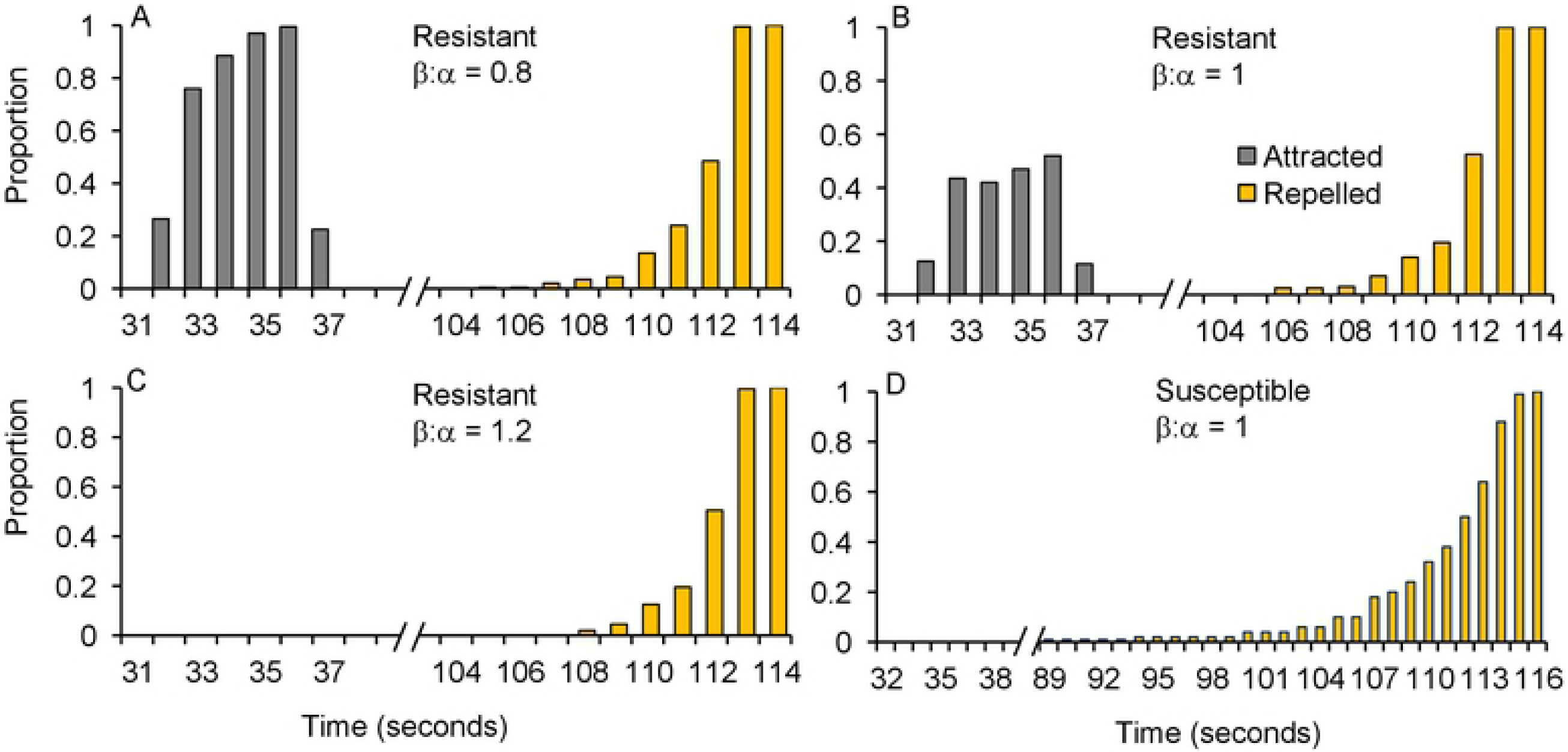
Frequency distribution of time when mosquitoes were attracted to the host or expelled from the simulation space, for non-resistant and resistant mosquitoes. Red bar: frequency distribution of resistant mosquitoes attracted to the odor release point at different times. Green bar: frequency distribution of resistant mosquitoes repelled from the simulation space at different times. Black curve: frequency distribution of non-resistant mosquitoes repelled from the simulation space at different times.

When β/α = 1, i.e., repellent and attractant were equally efficient, about half of the mosquitoes were able to reach the host before they were repelled (Figure 5B). On average these mosquitoes reached the host within 40.7 ± 3.7 s (range 32–48 s), and they were repelled within 275.9 ± 48.0 s. When β/α = 1.2, i.e., repellent was more efficient than attractant, none of the mosquitoes were able to reach the host and the average repel time was 277.6 ± 42.0 s (Figure 5C).

For the non-resistant mosquitoes, none reached the host before being repelled (Figure 5D). The distribution pattern of times when mosquitoes were repelled had a long, left-tailed repelling pattern (Figure 5D), which was rather different from resistant mosquitoes (Figures 4 & 5A-C). The first mosquito was repelled at 207 s, about 50 s faster than the resistant mosquitoes, and the proportion of repelled mosquitoes increased slowly from 207 s to 290 s (Figure 5D). The average repel time was 273.2 ± 16.1 s, very similar to the resistant mosquitoes.

## Discussion

The scale-up of malaria vector control using LLINs and IRS has led to widespread insecticide resistance as well as behavioral changes in *Anopheles* vectors. Vector resistance to pyrethroid insecticides and changes in biting and resting behavior pose serious challenges in malaria control. Spatial repellents are considered to be promising alternatives to the contact repellents currently in use, as they may delay or diminish the development of insecticide resistance by minimizing the intensity of selection pressure induced by contact-mediated toxicity mechanisms [37, 40]. However, since many spatial repellents contain the same or similar chemicals as those used in insecticides for current malaria control, their effectiveness can be compromised if insecticide-resistant mosquitoes behave differently when they encounter repellents, as has been demonstrated by recent field observations [51–58, 81, 84]. Our model simulation indicated that, in general, it took more time or a higher repellent concentration to expel mosquitoes with increased insecticide resistance. In addition, depending on resistance level and repellency strength, a proportion of mosquitoes continued to locate hosts even in the presence of a repellent. These results are similar to some field observations [51–58].

Resistant mosquitoes’ ability to reach the host even in the presence of repellent was as expected. This result is similar to findings from semi-field experiments for *Aedes aegypti* [84]. In their study on *Ae. aegypti*, Buhagiar and colleagues used a real house setting and they found that previously exposed *Ae. aegypti* were more likely to reach the host in a repellent-treated room [84]. In addition, if a mosquito in a repellent-treated room had not reached the host within 30 s, it never would [84]. This is similar to our simulation with non-resistant mosquitoes; i.e., none of the non-resistant mosquitoes reached the host in the spatial repellent–protected space, while resistant mosquitoes were able to reach the host before being expelled. As a result, higher concentrations of repellent were needed to deter blood-feeding by resistant mosquitoes [57]. In other words, physiological insecticide resistance may compromise the efficacy of spatial repellents. Here, repellency efficacy is important. Previous experiments showed that repellent may lose its efficacy against pyrethroid resistant mosquitoes [51–58]. Our simulation indicated that if relative repellency efficacy was lower compare to attractant efficacy, a high proportion of resistant mosquitoes were able to reach the host before they were repelled. In Buhagiar and colleagues experiments [85], they found 31% of the mosquitoes reached the host in the presence of repellent. When relative repellency efficacy was high, no mosquitoes could reach the host, indicating the importance of interactions between resistance and repellent.

Interestingly, in Buhagiar *et al* experiments [85], all *Aedes* mosquitoes in the control group reached the host at an average time of about 80 s, which was 50 s slower than treatment group. In Parker and colleagues experiments of insecticide sensitive *An. arabiensis* in Tanzania [68], they used human bait with no repellent in a hut and found out that mosquitoes first contacted the untreated net at a mean of 36 s after release, compared to 46 s in LLINs. In our simulations, all mosquito reached the host at an average of 83 s without repellent and mosquitoes might reach the host earlier when attractant concentration increased, more importantly, mosquitoes could reach the host in a short period of time (around 40 s) in the presence of repellent. These findings were in agreement with previous experiments [68]. The question is why do mosquitoes reach the host earlier in repellent protected space than non-repellent protected space? Previous studies indicated that insecticides/repellents increase mosquito activity, orthokinesis [86–88]. Kennedy’s study showed that mosquitoes get excited when they come in to contact with the repellent due to the poisoning effect [86]. When repellent is released in the space, mosquitoes sense the urgency to get a blood meal before they are repelled. This behavioral characteristics is often exhibited in the presence of repellents. An experimental study on cockroaches using DEET repellent showed that previously exposed insects exhibited an increased behavioral activity than non-exposed insects [89]. However, this behavioral characteristic needs to be tested on mosquitoes in carefully designed experiments.

A more interesting finding from this study is the slow repellency effect on non-resistant mosquitoes in an environment with both attractant and repellent (analogues to a room with a human host and spatial repellent). If no host were in the room, the same concentration of spatial repellent would repel non-resistant mosquitoes quickly. However, when a host was present, the repellency effect was significantly delayed, with a long left tail, as if the mosquitoes were resistant to repellent. This may be due to the mixture of attractant odors and repellent volatiles which confuses the mosquitoes as they are attracted by the human odor, and thus attempt to approach the host, but become disoriented by the repellent volatile chemicals. Similar finding was documented by Clark and Ray which indicated a prolonged activation of mosquito’s receptors in the presence of human odor (e.g. CO2) when in contact with airborne repellent compounds through mechanisms that are not well understood [90]. These results however need to be confirmed by semi-field or laboratory experiments.

There are differences between real-world and simulation settings. In typical African settings, houses usually have eaves which allow mosquitoes to exit, so the indoor use of spatial repellent may actually prevent mosquitoes from entering the house. However, the size of typical African houses are about 5m × 5m × 2.5 m, which is similar to the hut/house used in previous studies [68, 85]. Our simulation used an ideal enclosed space; mosquitoes were not allowed to escape. In a real-world setting, the respective diffusions of host odor and repellent volatiles may differ in many aspects. When burning coils in a room, one can smell the chemicals clearly at the beginning and see the smoke inside the room. Presumably, the concentration of repellent volatiles is higher than that of human odor, thus preventing mosquitoes from reaching the host. Studies indicate that spatial repellents compounds such as DEET, linalool, dehydrolinalool, catnip oil and citronella interfere with the attraction of mosquitoes to host odors by blocking natural responses to attractants, hence acting as attraction inhibitors and not repellents [88, 91, 92]. Results from Lucas *et al* study suggested that even in the presence of airborne pyrethroids, mosquitoes were able to detect host odors but were inhibited from feeding [93]. This effect is probably a result of pyrethroid – induced neural hyperexcitation, that can occur at much lower doses than those required for insect knockdown and mortality. In reality, after mosquitoes get a full blood meal, they will rest somewhere (such as the wall or ceiling in a typical African setting) to digest the meal. In the simulation study, we assumed mosquitoes did not move once they reached the host.

Theoretically, physiological resistance to insecticides may not necessarily affect how mosquitoes respond to repellents. However, selection experiments have found that spatial repellent–selected mosquitoes were insensitive to the spatial repellent [83]. Field studies found that field-collected pyrethroid-resistant *Ae. aegypti* were resistant to mosquito coils and other repellents [58,80]. In a malaria vector study in Kenya, Kawada and colleagues observed the lack of repellency effect of pyrethroids in the wild colony of *An. gambiae* s.s. with high resistance to pyrethroids, but not in the high-resistance wild colonies of *An. arabiensis* and *An. funestus* [52]. This was likely due to a difference in resistance mechanisms, since *An. arabiensis* and *An. funestus* are not affected by the knockdown effect. In this study, we assumed pyrethroid-resistant malaria vectors were insensitive to spatial repellent; however, the relationship between resistance and repellency needs further investigation.

In conclusion, malaria vector resistance to pyrethroid insecticides poses a serious challenge in malaria control. Spatial repellents have attracted significant attention in industry. Using SPP machine learning modeling, we simulated the potential impact of insecticide resistance on the response of malaria vectors to spatial repellents. We found that pyrethroid resistance may compromise the efficacy of spatial repellents, which warrants intensive investigation.

## Author Contributions

Conceived and designed the experiments: GZ, SK and GY. Performed the experiments: GZ, XW, DZ, MCL and GZ. Analyzed the data: GZ and MCL. Wrote the paper: GZ, SK and GY. Wrote the software used for the simulations: GZ and LY.

## Supplement

Figure S1. Illustrations of simulated 2D-projected mosquito flying paths. Scenarios: A) random walk; B) with attractant alone; C) with repellent and no insecticide resistance; D) with repellent and insecticide resistance; E) with repellent, attractant and resistance assuming β/α = 1.0. Simulation periods: blue colored section 0–10 s, green colored section 11–30 s and red colored section 31–60 s.

